# Association between alcohol consumption and Alzheimer’s disease: A Mendelian randomization Study

**DOI:** 10.1101/190165

**Authors:** Shea J. Andrews, Alison Goate, Kaarin J. Anstey

## Abstract

**INTRODUCTION:** Observational studies have suggested that light-moderate alcohol consumptions decreases the risk of Alzheimer’s disease, but it is unclear if this association is causal.

**METHODS:** Two-sample Mendelian randomization (MR) analysis was used to examine whether alcohol consumption, alcohol dependence or Alcohol Use Disorder Identification Test (AUDIT) scores were causally associated with the risk of Late Onset Alzheimer’s disease (LOAD) or Alzheimer’s disease age of onset survival (AAOS). Additionally, γ-glutamyltransferase levels were included as a positive control.

**RESULTS:** There was no evidence of a causal association between alcohol consumption, alcohol dependence or AUDIT and LOAD. Alcohol consumption was associated with an earlier AAOS and increased γ-glutamyltransferase blood concentrations. Alcohol dependence was associated with a delayed AAOS.

**DISCUSSION:** MR found robust evidence of a causal association between alcohol consumption and an earlier AAOS, but not alcohol intake and LOAD risk. The protective effect of alcohol dependence is potentially due to survivor bias.

**Research in Context:** *Systematic Review:* The authors reviewed the literature using online databases (e.g. PubMed). Previous research links light-moderate alcohol consumption to a decreased risk of Alzheimer’s disease (AD), however, prior studies based on observational study designs may be biased due to unmeasured confounders influencing both alcohol consumption and AD risk.

*Interpretation:* We used a two-sample Mendelian randomization (MR) approach to evaluated the causal relationship between alcohol intake and AD. MR uses genetic variants as proxies for environmental exposures to provide an estimate of the causal association between an intermediate exposure and a disease outcome. MR found evidence of a causal association between alcohol consumption and an earlier AD age of onset, suggesting that light-moderate alcohol consumption does not reduce risk of Alzheimer’s disease.

*Future Directions:* Future studies should use alterative study designs and account for additional confounders when evaluating the causal relationship between alcohol consumption and AD.

**Highlights:**

- We evaluated causal relationships between alcohol intake and Alzheimer’s disease
- Alcohol consumption is causally associated with an earlier Alzheimer’s age of onset
- No evidence of causal assocations between alcohol intake and Alzheimer’s risk

## 1. Introduction

The projected worldwide prevalence of dementia is expected to reach 74.7 million in 2030 [1]. This will have major implications for national health and social services, with the cost of caring for individuals living with dementia expected to rise from USD $818 billion in 2015 to USD $2 trillion in 2030 [1]. In the absence of any therapeutic interventions for dementia, successful intervention strategies that target modifiable risk factors to promote disease prevention are currently the only available approach that can have an impact on the projected rates of dementia.

Alcohol drinking is a modifiable behaviour that has emerged as a potential protective factor for dementia. A recent overview of systematic reviews that investigated the association between alcohol consumption and dementia or cognitive decline, identified three high quality systematic reviews that meet their inclusion criteria [2]. Two of the three systematic reviews conducted a meta-analysis and concluded that light-moderate alcohol consumption in contrast to abstainers was correlated with a 25-38% reduction in risk of Alzheimer’s disease (AD), vascular dementia (VaD) and all cause dementia (ACD) [3,4]. In contrast the third review, performed a qualitative analysis and concluded that the literature did not provide concrete evidence of a causal association between alcohol consumption and AD [5]. A more recent meta-analysis found light-moderate alcohol consumption to be protective [6]. Light alcohol consumption corresponding to either 4 drinks/week, 6g/day or 1 times/week provided the greatest risk reduction in ACD, while excessive alcohol consumption of 23 drinks/week or 12.5g/day significantly increased risk of ACD. Furthermore, qualitative analysis indicated that the protective effects only existed for wine consumption [6]. Observational studies however may underestimate the detrimental effects of long-term excessive alcohol consumption, which is associated with the development of alcohol-related dementia (ARD), due to participants with ARD not being recruited or lost to follow-up [7].

Several mechanisms have been proposed that may underlie the observed protective effects of moderate alcohol consumption on dementia. First, moderate alcohol consumption has been associated with increased levels of circulating high-density lipoprotein cholesterol, apolipoprotein AI and adiponectin and decreased fibrinogen levels [8]. The cardio protective effects of these biomarkers decrease cardiovascular disease risk, thus reducing the risk of cerebrovascular injury. Second, moderate alcohol consumption is associated with an anti-inflammatory effect that could moderate the neuroinflammatory response observed in Alzheimer’s disease [9,10]. Finally, resveratrol, found in red wine, is proposed to have both neuroprotective effects and anti-oxidant properties that reduce neuronal cell death [11].

While accumulating evidence suggests that light-moderate alcohol consumption is associated with a reduced risk of dementia, there are several limitations within the literature that temper the positive interpretation of these results. First, ascertainment criteria may bias result, with heavy drinkers who already show signs of cognitive impairment screened out of studies [3]. Second, alcohol consumption is associated with an increased mortality risk, potentially resulting in participants being lost to follow-up, prior to a dementia diagnosis. This would result in an underestimate of dementia incidence and bias estimates of relative risk [12]. Third, the abstainer comparison groups within studies are often composed of life-time abstainers and former drinkers who may have reduced or quit drinking because of other detrimental health outcomes [13,14]. As such it is possible that confounding health factors increase dementia risk in non-drinkers. Fourth, the apparent protective association between alcohol consumption and dementia may be confounded by other health/lifestyle characteristics, such as lower cardiovascular risk factors, that are associated with a reduced incidence of dementia [3,4]. In particular, socioeconomic status and prior intelligence both influence the amount and type of alcohol consumed, and as such may play an important confounding role in the alcohol-dementia relationship [15].

As such, the current observational studies are limited by issues of confounding and reverse causality. In the absence of randomized control trials, a novel method for estimating causal effects of risk factors in observational studies using genetic variants is Mendelian randomization (MR).

Mendelian randomization uses genetic variants as proxies for environmental exposures to provide an estimate of the causal association between an intermediate exposure and a disease outcome (Figure 1) [16]. MR is similar to a ‘genetic randomized control trial’ due to the random allocation of genotypes from parents to offspring and are thus not affected by reverse causation and are independent of confounding factors that may influence disease outcomes (Figure 1) [16]. The genetic variants used in MR act as an instrumental variable (IV) and if the assumptions hold for the genetic variant (Figure 3), any association between the genetic variants and the disease outcome must come via the variant’s association with the exposure. This implies that the exposure is causally related to the outcome.

**Figure 1:**
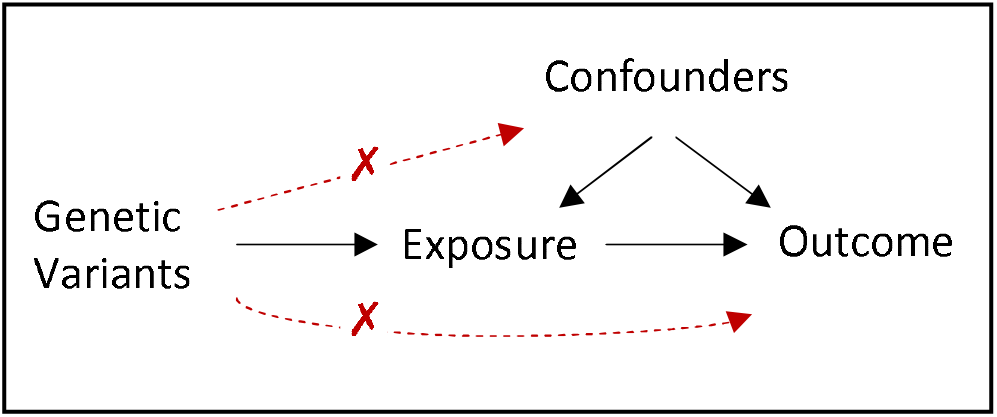
Model for a Mendelian randomization study. Genetic variants known to be associated with the exposure (Non-zero effect assumption) are used to estimate if the exposure causally influences the outcome. The genetic variable is assumed not to be associated with confounders (Independence assumption) or the outcome (exclusion restriction assumption).

Mendelian randomization has been used to investigate the causal effects of potential risk factors on the risk of AD. Genetically predicted higher systolic blood pressure, greater daily smoking quantity and increased adult height were causally associated with a lower risk of AD, genetically decreased vitamin D and plasma apolipoprotein E, levels were causally related to increased risk of AD [17–19]. Conversely, MR did not support evidence of causal links between glycemic traits, type 2 diabetes, BMI, educational attainment, leptin concentrations, circulating homocysteine levels or coffee consumption and risk of AD [17–19].

In this study, we perform a two-sample MR analysis [20]to assess the causal effect of alcohol consumption, alcohol dependence and AUDIT scores on the risk of developing Alzheimer’s disease and Alzheimer’s disease age of onset survival. Additionally, we also perform a two-sample MR analysis with alcohol consumption and γ-glutamyl transferase (GGT) as a positive control, where the causal effects between GGT and alcohol intake have been strongly established [21].

## 2. Methods

### 2.1 Instruments

GWAS were obtained for each exposure of interest and single nucleotide polymorphisms (SNPs) associated with each trait at genome wide significance (*P* < 5 × 10^−8^) were extracted. SNPs were clumped to obtain independent loci using a threshold of linkage disequilibrium (LD) r^2^ > 0.001 and a distance of 1000kb in PLINK [22]. SNPs associated with the exposure were extracted from each outcome GWAS and where SNPs for the exposure were not available in the outcome GWAS PLINK was used to identify proxy SNPs that were in LD (r^2^ > 0.8; 1000 Genomes European reference population). The exposure– outcome datasets were harmonized using the TwoSampleMR package, allowing forward strand ambiguous SNPs to be inferred using allele frequency information, with strand ambiguous SNPs with intermediate allele frequencies (AF > 0.42) removed from analysis [23]. The proportion of variance in the phenotype explained by each instrument and F-statistic were calculated as previously described [24,25].

#### Alcohol Consumption

SNPs associated with alcohol consumption were selected from a meta-analysis of GWAS in 25 cohorts performed by the GWAS & Sequencing Consortium of Alcohol and Nicotine use (GSCAN) consortium in 537,349 participants of European ancestry [26]. Alcohol consumption was scored as the number of self-reported drinks per week and was left anchored to 1 and log transformed. The genomic control inflation factor was 1.24, and statistical models adjusting for age, age^2^, sex, genetic principal components and other study-specific covariates. Forty four SNPs were extracted, with an F-statistic of 70.8 and explaining 0.58% of the total variance.

#### Alcohol Dependence

SNPs associated with Alcohol dependence were selected from a meta-analysis of GWAS in 28 cohorts performed by the Psychiatric Genomic Consortium (PGC) in 46,568 (11,569 cases and 34,999 controls) individuals of European ancestry [27] with 51.6% of the sample female. Alcohol dependence was diagnosed using clinician rating or semi-structured interviews following DSM-IV criteria, with statistical models adjusted for sex and genetic principal components. Twenty SNPs were extracted, with an F-statistic of 24.6, explaining 1.14% of the total variance.

#### Alcohol Use Disorder Identification Test (AUDIT)

SNPs associated with AUDIT scores were obtained from a GWAS in 121,604 individuals of European ancestry from the UK Biobank Cohort [28]. The sample was 56.2% female and the mean age was 56.1 (SD 7.7) years. AUDIT was scored as the total sum of items 1-10 and log10 transformed to approximate a normal distribution, with statistical models adjusted for age, sex, genotyping array and genetic principal components. As allele frequencies were not reported, Haplotype Reference Consortium [29] or EUR 1000 Genomes [30] allele frequencies were used. Eleven SNPs were extracted, with an F-statistic of 59.3, explaining 0.53% of the variance.

#### Late Onset Alzheimer’s Disease (LOAD)

SNPs associated with LOAD were obtained from a meta-analysis of 4 previously published GWAS datasets: the European Alzheimer’s Disease Initiative (EADI), the Alzheimer Disease Genetics Consortium (ADGC), Cohorts for Heart and Aging Research in Genomic Epidemiology (CHARGE), and Genetic and Environmental Risk in AD (GERAD) and includes a sample of 17,008 LOAD cases and 37,154 cognitively normal elderly controls [31]. Participants in IGAP were of European ancestry, the average age was 71 years and 58.4% of participants were women. The EUR 1000 Genomes Project population reference was used to impute genotypes, the genomic control inflation factor was 1.087, and analysis was adjusted for age, sex and ancestry using principal component scores. As allele frequencies were not reported, allele frequencies from a GWAS of Alzheimer’s age of onset survival, based on a subset of the IGAP cohort, were used [32].

#### Alzheimer’s disease Age of Onset Survival (AAOS)

SNPs associated with AAOS were obtained from a genome-wide survival analysis of 14,406 AD cases and 25,849 controls from IGAP [32]. Participants were of European ancestry, had an average age of 77.49 years (SD = 8.4) and 60.35% were women. Statistical analysis consisted of a genome-wide Cox proportional hazards model using age at onset for cases and age at last assessment for controls, adjusted for sex, site and principal components.

#### γ-glutamyltransferase (GGT)

SNPs associated with γ-glutamyltransferase blood concentration (log_10_ transformed IU/L) were obtained from a GWAS performed in 61,089 participants of European ancestry, with an average age of 52.8 years and 50.4% women [33]. HapMap build 36 and dbSNP build 126 were used to impute missing genotypes, the genomic control inflation factor was 1.005, and analysis was adjusted for age, sex, case/control status, and ancestry using principal component scores.

### 2.2 Mendelian Randomization Analysis

All statistical analyses were conducted using R version 3.5.2 [34], with the MR analysis performed using the ‘TwoSampleMR’ package [23].

The SNP-exposure and SNP-outcome coefficients were combined in a fixed-effects meta-analysis using an inverse-variance weighted approach to give an overall estimate of causal effect [20]. This is equivalent to a weighted regression of the SNP-outcome coefficients on the SNP-exposure coefficients with the intercept constrained to zero. This method assumes that all variants are valid instrumental variables based on the MR assumptions (Figure 1). In order to account for potential violations of the assumptions underlying the IVW MR analysis, we compared the IVW results to other methods known to be more robust to horizontal pleiotropy, but at the cost of reduced statistical power. First, a weighted median MR was performed, which allows for 50% of the instrumental variables to be invalid [35]. Second, MR-Egger regression was performed, which allows all the instrumental variables to be subject to direct effects (i.e. horizontal pleiotropy) [36], with the intercept representing bias in the causal estimate due to pleiotropy and the slope representing the causal estimate. The causal estimate of the IVW analysis expresses the causal increase in the outcome (or log odds of the outcome for a binary outcome) per unit change in the exposure.

Mendelian randomization pleiotropy residual sum and outlier (MR-PRESSO) test was also used to detect and correct for horizontal pleiotropic outliers [37]. MR-PRESSO conducts a global test of heterogeneity to detect horizontal pleiotropy by regressing the SNP-outcome associations on the SNP-exposure associations and comparing the observed distance of each SNP from the regression line with the distance expected under the null hypothesis of no pleiotropy. In the case of horizontal pleiotropy, the MR-PRESSO outlier test compares individual variants expected and observed distributions to identify outlier variants. Where significant outliers are detected, they were removed from the analysis to obtain an unbiased causal estimate.

Power calculations were conducted using the mRnd power calculation tool [38]. Given a sample size of 54,162 with the proportion of cases equal to 0.314 (IGAP), this study was adequately powered to detect an OR of any AD of 0.966 for alcohol consumption.

## 3. Results

Harmonized SNP exposure–outcome datasets and SNPs indicated as outliers are available in Supplementary Table 1. The causal estimates for the IVW, MR-Egger and Weighted median analysis, number of SNPs and number of outliers removed for each exposure – outcome pair are presented in Table 1. Causal estimates prior to and after outlier removal are presented in Supplementary Table 2. Figure 2 shows scatter plots of the SNP-exposure and SNP-outcome association estimates and their corresponding MR causal estimates.

**Table 1:** IVW MR estimates after outlier removal for the causal associations of alcohol consumption, alcohol dependence and AUDIT scores with Alzheimer’s disease, Alzheimer’s age of onset and *γ-glutamyltransferase*.

| Exposure | SNPs | outliers | IVW | MR-Egger | Weighted Median | MR-PRESSO | MR-Egger |
| --- | --- | --- | --- | --- | --- | --- | --- |
|  | *N* | *N* | b (se) | b (se) | b (se) | Global<br>*p* | Intercept<br>*p* |
| **<u>Late onset Alzheimer's disease</u>** |  |  |  |  |  |  |  |
| Alcohol Consumption | 41 | 1 | -0.04 (0.13) | -0.21 (0.22) | -0.21 (0.19) | 0.0578 | 0.286 |
| Alcohol Dependence | 17 | 0 | -0.02 (0.03) | -0.04 (0.06) | -0.05 (0.04) | 0.275 | 0.652 |
| AUDIT - Total | 7 | 1 | -0.79 (0.69) | -1.47 (2.23) | -0.96 (0.98) | 0.017 | 0.732 |
| **<u>Alzheimer's age of onset survival</u>** |  |  |  |  |  |  |  |
| Alcohol Consumption | 42 | 2 | 0.7 (0.18)*** | 0.6 (0.55) | 0.72 (0.26)** | 0.9152 | 0.844 |
| Alcohol Dependence | 19 | 1 | -0.07 (0.03)* | -0.22 (0.09)* | -0.08 (0.05). | 0.033 | 0.079 |
| AUDIT - Total | 9 | 0 | -0.73 (0.66) | -2.09 (1.14) | -0.48 (0.92) | 0.307 | 0.185 |
| **<u>γ-glutamyltransferase</u>** |  |  |  |  |  |  |  |
| Alcohol Consumption | 32 | 2 | 0.1 (0.03)** | 0.15 (0.1) | 0.12 (0.05)* | 0.5558 | 0.553 |
| Alcohol Dependence | 11 | 0 | -0.01 (0.01) | -0.05 (0.03) | -0.01 (0.01) | 0.091 | 0.197 |
| AUDIT - Total | 4 | 2 | 0.26 (0.18) | - | - | - | - |

\* $p < 0.05$ ; \*\* $p < 0.01$ ; \*\*\* $p < 0.001$

**Figure 2:**
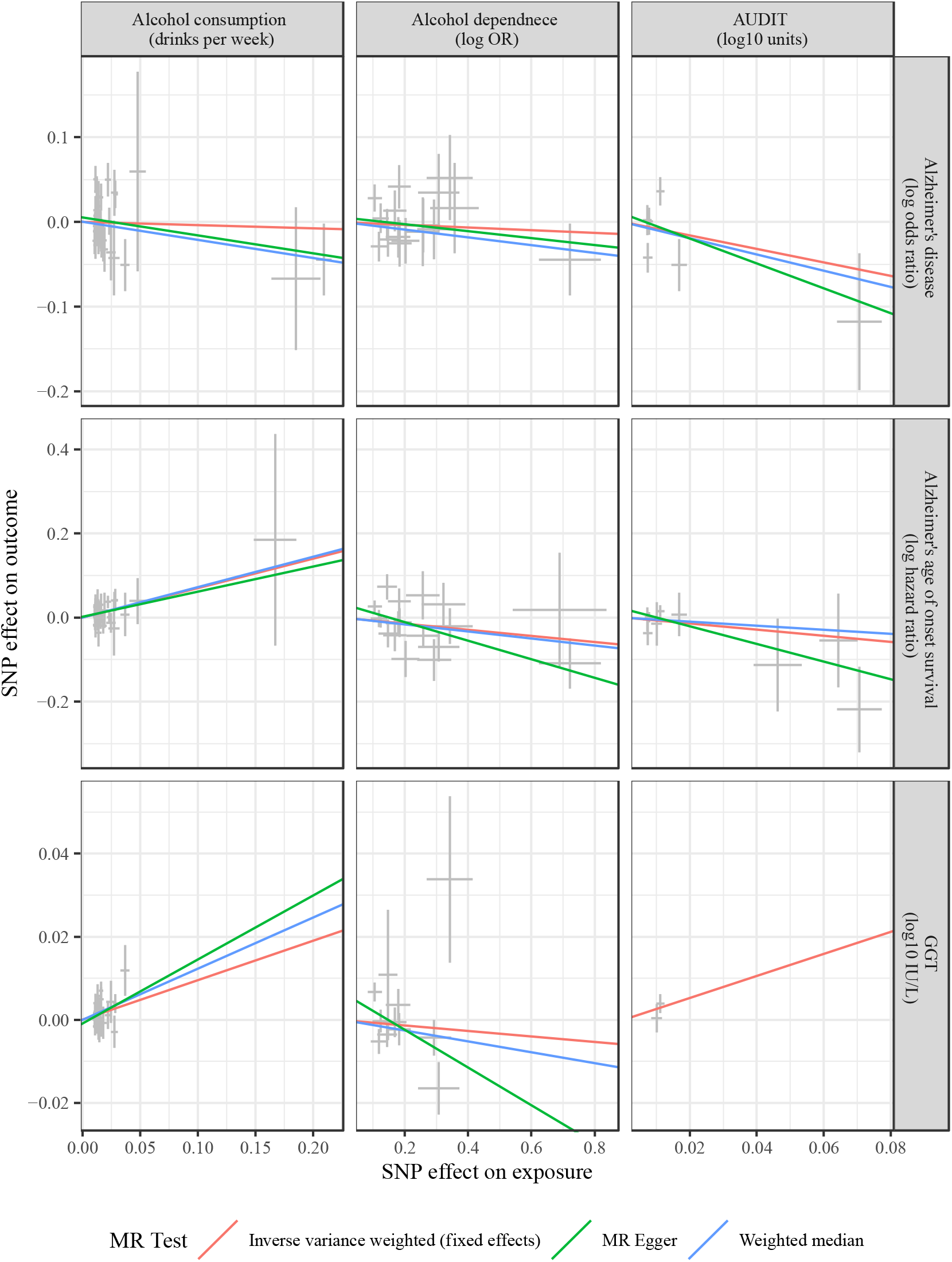
Scatter plots of the SNP effects on the exposure and the outcome. The SNP-exposures association estimates have been recoded in order to make the SNP-exposure associations positive to facilitate interpretation. Regression lines represent the causal effect of the exposure on the outcome using IVW, MR-Egger and Weighted median to estimate the causal effects.

### 3.1 Late Onset Alzheimer’s Disease

Using IVW, genetically predicted alcohol consumption (OR [95%CI] = 0.96 [0.74, 1.25], p = 0.775), alcohol dependence (OR [95%CI] = 0.98 [0.93, 1.04], p = 0.588) and AUDIT scores (OR [95%CI] = 0.45 [0.12, 1.75], p = 0.25) were not causally associated with LOAD.

### 3.2 Alzheimer’s Age of Onset Survival

In the IVW analysis, genetically predicted alcohol consumption was associated with an earlier AAOS (HR [95%CI] = 2.02 [1.42, 2.87], p = 9.4 × 10^−05^). In the corresponding MR-PRESSO Global Test and MR-Egger Intercept test, there was no evidence of heterogeneity or pleiotropy. Additionally, the Weighted Median analysis was significant, and the MR-Egger results trended in the same direction. After outlier removal, genetically predicted alcohol dependence was associated with a delayed AAOS (HR [95%CI] = 0.93 [0.87, 0.99], p = 0.031). In the sensitivity analysis, the MR-Egger analysis was also associated with a delayed age of onset and there was no evidence that the intercept differed from the null. Genetically predicted AUDIT scores were not significantly associated with AAOS.

### 3.3 γ-glutamyltransferase

In the IVW analysis, after outlier removal genetically predicated alcohol consumption was associated with increased GGT blood concentrations (β [SE] = 0.1 [0.03], p = 0.003) In the sensitivity analysis there was no evidence of heterogeneity or pleiotropy based on the MR-PRESSO Global Test and MR-Egger Intercept test and the Weighted Median analysis was also significant. Genetically predicted alcohol dependence and AUDIT scores were not associated with GGT blood concentrations.

## 4. Discussion

This is the first Mendelian randomization study to examine the causal association between alcohol intake and Alzheimer’s disease. Genetically predicted alcohol consumption was not associated with the risk of developing Alzheimer’s disease, but was observed to be associated with an earlier Alzheimer’s age of onset, such that individuels with 1SD higher consumption of alcohol are twice as likely to develop Alzheiemr’s at a given point of time. resulting in a 66% probability of an earlier age of onset. This finding was robust to pleiotropy and heterogeneity after outlier removal, and the causal estimates from the Weighted median and MR-Egger sensitivity analyses were concordant with the IVW analysis suggesting that the causal effect was robust to violations of the underlying MR assumptions. Additionally, alcohol consumption was associated with levels of γ-glutamyltransferase, confirming that the instrumental variable was a suitable proxy for alcohol consumption in the MR analysis. This suggests that light-moderate alcohol intake is unlikely to reduce the risk of Alzheimer’s disease but is associated with an earlier age of onset.

This finding contradicts those from recent meta-analyses and systematic reviews [2,6], which have suggested that light-moderate alcohol consumption is protective against dementia. However, all prior findings have been conducted in observational studies that assume no confounders influence the reported results and are limited by selection bias, an underlying illness-death structure, and the heterogeneous nature of the abstainer comparison group. Like randomized control trials, MR analyses reduce confounding and reverse causality due to the random allocation of genotypes from parents to offspring. This can allow for a more robust inference of causal effects. As such the results from this study provide further support for the cautious interpretation of the proposed cognitive health benefits of alcohol [14], and further highlights that future observational studies need to account for potential confounding factors.

Alcohol dependence was also observed to be associated with a delayed Alzheimer’s age of onset, such that at a given point of time individules with alcohol dependence are 7% less likely to develop AD. Furthermore, the results from the sensitivity analysis were concordant and significant, suggesting that the causal effect was robust to violations of the underlying MR assumptions. Nevertheless, this result contradicts the observational literature, where excessive alcohol consumption (in reference to light-moderate consumption) defined as more than 14 drinks/week increases the risk of dementia in a linear fashion [39]. However, alcohol dependence is also associated with a higher risk of somatic diseases (e.g. cancer, cardiovascular, metablic disease) and mortality [40]. As such, these contradictory results may be due to survivor bias, whereby mortality due to competing risks affects selection into the target study, thereby biasing causal estimates. In particular in MR studies conducted in elderly populations, if the genetic variants associated with the exposure increase mortality rates, individuals with those genetic variants who are still alive at the onset of the study are also less likely to have other risk factors associated with mortality [41–43]. This means that the genetic variants will be associated with other risk factors that affect the outcome and thus violate the assumptions underlying MR [41–43]. The bias introduced by survival effects is large for exposures that strongly affect survival, however, when selection effects are weak or moderate, selection bias does not adversely affect causal estimates [43]. Surivor bias may also explain the contradictory findings observed in this MR analysis, where alchol consumption being associated with a earlier AAOS and alchol depence being associated with a delayed AAOS. Additionally, no association between alcohol dependence or AUDIT scores and GGT was observed, suggesting that the instruments may not have being suitable proxies for the MR analysis.

The results from this study should be interpreted in conjunction with some limitations. First, we cannot be certain the selected SNPs do not violate the exclusion-restriction assumption. However, we did use MR-PRESSO and MR-Egger regression to estimate the extent to which heterogeneity and pleiotropy may bias the reported results, and have reported those results that are robust to the violations of MRs assumptions. Second, given the use of separate samples, we were unable to test whether the association with AD varied by level of alcohol consumption or by other covariates such as age or gender. Given the proposed dose-response relationship between alcohol consumption and dementia, a causal relationship could be expected between excessive alcohol consumption and increased risk of dementia. Nevertheless, the use of alcohol dependence as an instrumental variable should address this issue. Third, in two-sample MR studies it is assumed that both samples come from comparable populations, and confounding due to population stratification cannot be ruled out [44]. However, in this study, the corresponding GWAS were all conducted in samples of European ancestry and genomic control for ethnicity was conducted in each individual GWAS. However, this does limit the interpretation of these results to other ethnic groups. Fourth, non-random selection into the analytical cohorts [44], particularly in the UK Biobank datasets where the response rate for recruitment was low and individuals had higher average levels of educational attainment and general health. [45] may bias results. Finally, canalization whereby the genetic effect of alcohol consumption on AD is modified via compensatory mechanisms may attenuate the association of genetically determined alcohol intake with AD [46]. For example upregulation of *ALDH2* gene expression in response to excessive alcohol consumption.

Despite these limitations, this study has a number of strengths. First, this study takes advantage of publicly available datasets to gain more precise estimates and greater statistical power due to the large sample sizes associated with the GWAS for alcohol consumption (*n* = 537,349), AUDIT scores (*n* = 121,604) alcohol dependence (*n* = 46,568), and Alzheimer’s disease (*n* = 54,162). This study was adequately powered to detect a significant association between a 1-SD increase in units of alcohol consumed per week on AD risk of at least OR = 0.79 or an OR = 1.28. Second, MR is less prone to the bias of observational studies, particularly in relation to reverse causation and confounding, providing a more robust estimate of causal relationships between exposures and outcomes.

In conclusion, this Mendelian randomization study did not find evidence of a causal relationship between alcohol intake and risk for Alzheimer’s disease, but did find evidence that alcohol consumption was associated with earlier Alzheimer’s disease age of onset. This contrasts with recent findings from systematic reviews which have suggested that light-moderate alcohol consumption is associated with reduced risk of dementia outcomes. This discrepancy could be due to limitations in the methodology of observational studies including, selection effects, study attrition, the heterogeneous nature of abstainer groups or confounding. As such our results reiterate that abstainers should not initiate alcohol consumption to improve ‘cognitive health’ and further suggest that alcohol consumption should be reduced considering the overall harmful effects of alcohol.

## Acknowledgments

KJA is funded by NHMRC Research Fellowship No. 1002560. S.J.A. was suported by the ARC Centre of Excellence in Population Ageing Research, ARC grant CE1101029 and the JPB Foundation (http://www.jpbfoundation.org). AMG was supported by the National Institute on Alcohol Abuse and Alcoholism (U10AA08401) and the JPB foundation.

This work was made possible by the generous sharing of GWAS summary statistics. We thank Professor John Chambers for the provision of the summary results data for the GWAS on liver enzyme concentrations in plasma. We thank the International Genomics of Alzheimer’s Project (IGAP) for providing summary results data for these analyses. The investigators within IGAP contributed to the design and implementation of IGAP and/or provided data but did not participate in analysis or writing of this report. IGAP was made possible by the generous participation of the control subjects, the patients, and their families. The i–Select chips was funded by the French National Foundation on Alzheimer’s disease and related disorders. EADI was supported by the LABEX (laboratory of excellence program investment for the future) DISTALZ grant, Inserm, Institut Pasteur de Lille, Université de Lille 2 and the Lille University Hospital. GERAD was supported by the Medical Research Council (Grant n° 503480), Alzheimer’s Research UK (Grant n° 503176), the Wellcome Trust (Grant n° 082604/2/07/Z) and German Federal Ministry of Education and Research (BMBF): Competence Network Dementia (CND) grant n° 01GI0102, 01GI0711, 01GI0420. CHARGE was partly supported by the NIH/NIA grant R01 AG033193 and the NIA AG081220 and AGES contract N01–AG–12100, the NHLBI grant R01 HL105756, the Icelandic Heart Association, and the Erasmus Medical Center and Erasmus University. ADGC was supported by the NIH/NIA grants: U01 AG032984, U24 AG021886, U01 AG016976, and the Alzheimer’s Association grant ADGC–10–196728. IGAP URL: http://web.pasteur-lille.fr/en/recherche/u744/igap/igap_download.php

